# RNA- and ATAC-sequencing Reveals a Unique *CD83+* Microglial Population Focally Depleted in Parkinson’s Disease

**DOI:** 10.1101/2023.05.17.540842

**Authors:** Z.K. Chatila, A. Yadav, J. Mares, X. Flowers, T.D. Yun, M. Rashid, R. Talcoff, Z. Pelly, Ya Zhang, P.L. De Jager, A. Teich, R. Costa, E. Area Gomez, G. Martins, R. Alcalay, J.P. Vonsattel, V. Menon, E.M. Bradshaw, S. Przedborski

## Abstract

All brain areas affected in Parkinson’s disease (PD) show an abundance of microglia with an activated morphology together with increased expression of pro-inflammatory cytokines, suggesting that neuroinflammation may contribute to the neurodegenerative process in this common and incurable disorder. We applied a single nucleus RNA- and ATAC-sequencing approach using the 10x Genomics Chromium platform to postmortem PD samples to investigate microglial heterogeneity in PD. We created a multiomic dataset using substantia nigra (SN) tissues from 19 PD donors and 14 non-PD controls (NPCs), as well as three other brain regions from the PD donors which are differentially affected in this disease: the ventral tegmental area (VTA), substantia inominata (SI), and hypothalamus (HypoTs). We identified thirteen microglial subpopulations within these tissues as well as a perivascular macrophage and a monocyte population, of which we characterized the transcriptional and chromatin repertoires. Using this data, we investigated whether these microglial subpopulations have any association with PD and whether they have regional specificity. We uncovered several changes in microglial subpopulations in PD, which appear to parallel the magnitude of neurodegeneration across these four selected brain regions. Specifically, we identified that inflammatory microglia in PD are more prevalent in the SN and differentially express PD-associated markers. Our analysis revealed the depletion of a *CD83* and *HIF1A-*expressing microglial subpopulation, specifically in the SN in PD, that has a unique chromatin signature compared to other microglial subpopulations. Interestingly, this microglial subpopulation has regional specificity to the brainstem in non-disease tissues. Furthermore, it is highly enriched for transcripts of proteins involved in antigen presentation and heat-shock proteins, and its depletion in the PD SN may have implications for neuronal vulnerability in disease.

## Introduction

Parkinson’s disease (PD) is the second most common neurodegenerative disorder after Alzheimer’s Disease (AD)^1^. PD, like other neurodegenerative disorders, is characterized by the dysfunction and death of specific subsets of neurons^2^. Indeed, in PD, even within affected areas striking differential neuronal susceptibility is observed^3^. Ventral midbrain dopaminergic DA neurons in the substantia nigra pars compacta (SN) are consistently more affected than those in the ventral tegmental area (VTA)^1, 2^. Thus, while research efforts have focused on identifying drivers of neuronal death in PD^4^, elucidating determinants of neuronal resilience in PD may have far-reaching implications for both our understanding of PD pathobiology and our ability to develop effective disease-modifying therapies for this disabling, incurable disease.

Traditionally, neuronal susceptibility has been studied from a cell-autonomous perspective, but mounting evidence suggests that CNS-resident and peripheral immune cells also contribute to neurodegeneration^5^. The idea that PD pathogenesis may involve an immune component emanates in part from observations at the sites of neuropathology, including accumulation of microglia with an activated morphology, increased expression of pro-inflammatory cytokines, and the presence of T cells^6–9^. Moreover, genes implicated in the regulation of leukocyte/lymphocyte activity and cytokine-mediated signaling^10^ as well as genetic variants for the microglial receptor *TREM2* have been linked to increased risk for several neurodegenerative disorders including PD^11^. Furthermore, PD susceptibility alleles are enriched in genes with microglial-specific cis-regulatory effects^12, 13^. Epidemiological studies in humans with parkinsonism and studies in experimental models of PD further support a role for neuroinflammation in its pathogenesis^14–23^.

Importantly, microglia are now appreciated to be highly heterogenous and plastic cells that modulate their gene expression and function in response to the CNS microenvironment^24–26^. We posit that variations in microglial responses contribute to the differential neuronal susceptibility observed in PD. We therefore sought to investigate microglial heterogeneity at the single-nucleus level in selected post-mortem human brain regions from PD and non-Parkinson’s disease controls (NPCs). In this work, we have identified alterations in microglial subpopulations in PD that support the theory of a gain in proinflammatory microglia and loss of beneficial microglial in disease^27, 28^. We have characterized proinflammatory and proliferative microglial subpopulations that are enriched in the SN from PD donors. Conversely, we discovered an activated microglial population that is present in the SN of NPCs and in less-affected regions in PD but depleted in the PD SN. This microglial subpopulation is enriched for gene sets involved in antigen presentation and chaperone proteins and appears to be unique to the brainstem, hypothalamus, and cerebellum in the non-diseased brain. This microglial subpopulation may therefore be poised to maintain protective neuro-immune dynamics in regions including the brainstem, and its loss may have implications for selective neuronal vulnerability in PD.

## Results

### Building an Unbiased Transcriptomic Dataset from Human PD Brain Tissues

To advance our understanding of the role of innate immune cells in PD, we sought to capture the cellular and molecular heterogeneity of the local CNS myeloid cell response associated with this neurodegenerative disorder. Accordingly, we performed single-nucleus multiome RNA-sequencing (snRNA-seq) and assay for transposase-accessible chromatin with sequencing (ATAC-seq) on fresh frozen substantia nigra pars compacta (SN) tissues from 19 individuals with idiopathic PD and 14 aged-matched NPCs. This midbrain region was selected as it represents a prototypic brain area that is most affected in PD^29^. To acquire additional insights into the myeloid response in PD, we sought to capture the immune cell dynamics across regions that have different degrees of neuronal loss in this disease (Extended Data Fig. 1)^30–35^. Accordingly, in addition to the SN, we also sequenced nuclei from three other brain regions from the same cohort of PD donors, the ventral tegmental area (VTA), the substantia inominata (SI), and the rostral hypothalamus (HypoTs). These regions provide a range of neurodegeneration, with the greatest degree of neuronal loss in the SN, followed by the VTA, the SI, and the lowest degree of neuronal loss in the HypoTs (Extended Data Fig. 1). Table 1, Supplementary Table 1, and Supplementary Table 2 summarize the clinical and neuropathological information for both PD and NPC groups.

**Table 1:**
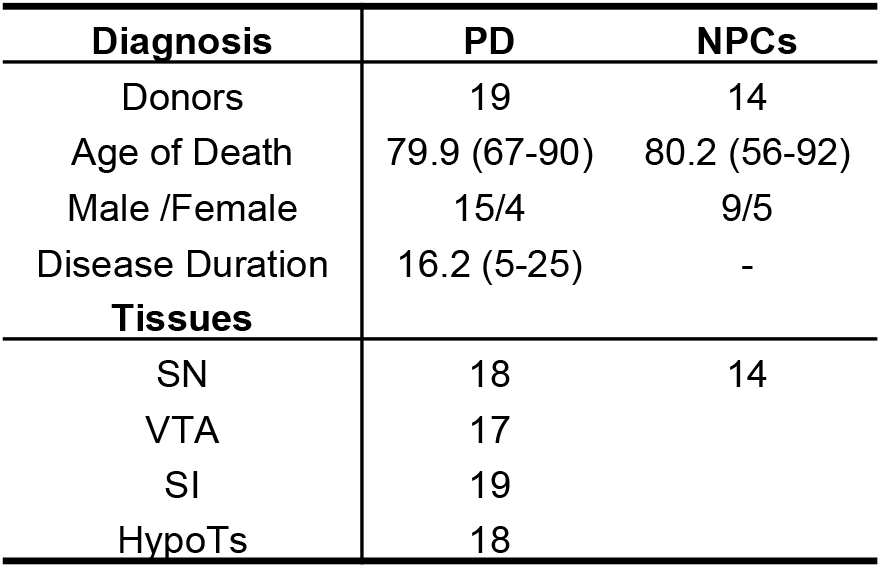
Summary of Parkinson Disease (PD) and non-PD control (NPC) Donors and Tissue Samples Included in MultiomeSequencing.

**Figure 1.**
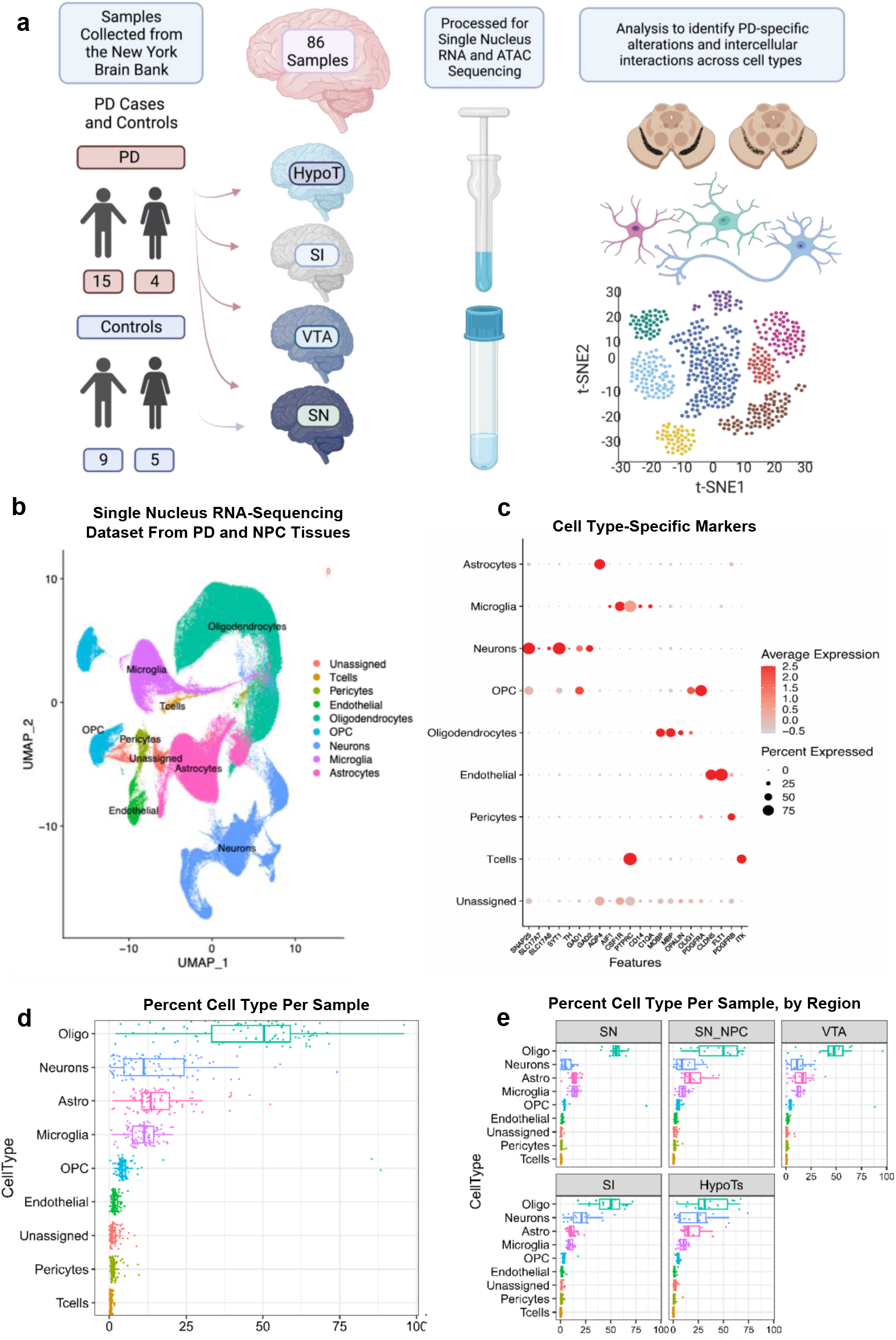
Unsupervised clustering analysis of snRNA-seq. **a.** Fresh frozen brain tissues for single nucleus multiome RNA- and ATAC-seq from 19 individuals with Parkinson’s disease (PD) and 14 non-PD controls (NPCs). NPC tissue samples include the substantia nigra (SN), and PD tissue samples include the SN, ventral tegmental area (VTA), substantia inominata (SI), and hypothalamus (HypoTs). **b.** UMAP plot from snRNA-seq showing the major cell classes present in the tissue samples sequenced from PD and NPC donors. Each dot represents a nucleus, and dots are color coded to indicate cell class grouping. **c.** Gene expression patterns of major cell classes for cell-type specific markers. Each column represents a selected key gene, and each row represents the indicated major cell class. **d**. Box plot showing the proportion of each cell type per sample in the dataset, across all regions. Each dot represents one tissue sample. **e.** Box plot showing the proportion of each cell type per sample in the dataset, split by region of origin. Each dot represents one tissue sample.

This snRNA- and ATAC-sequencing workflow from these PD and NPC samples (outlined in Fig. 1a) generated a median of 5,040 unique molecular identifiers (UMIs) with 2,058 median genes per nucleus, a median of 3,805 open chromatin peaks detected per nucleus, and a median of 4,812 nuclei per sample after quality control (see Methods). These values indicate that the quality of our single-nucleus profiling is consistent with previous transcriptomic studies on post-mortem subcortical tissue^4, 36^.

We began our analysis by integrating the snRNA-seq datasets from all PD and NPC tissue samples and performed unsupervised clustering analysis of this integrated dataset to identify major cell classes (Fig. 1b). We used gene expression of canonical markers to categorize nuclei (Fig. 1c) into neurons (*SNAP25, SLC17A7, SLC17A6, SYT1, TH, GAD1, GAD2*), oligodendrocytes (*MOBP*, *MBP, OPALIN*), oligodendrocyte precursor cells (*OLIG1*, *PDGFRA*), astrocytes (*AQP4*), microglia (*AIF1, P2RY12, CSF1R, C1QA PTPRC, CD14*), T cells (*CD3, ITK*), vascular cells (*CLDN5, RGS5*), endothelial cells (*FLT1, CD13*), and pericytes (*PDGFRB*).

Oligodendrocytes account for the largest population of nuclei in each tissue, on average comprising 45.6% of nuclei per sample. Neurons and astrocytes have similar frequencies and comprise an average of 15.3% and 16.1% of nuclei per sample, respectively. Microglia account for an average of 11.1% of nuclei per sample. Other cell types account for an average of <10% of all nuclei per sample, including oligodendrocyte precursor cells (6.4%), endothelial cells (1.9%), pericytes (1.5%), and T cells (0.5 %) (Fig. 1d and 1e). Although these proportions may not represent cell type composition in intact tissue, these results indicate that we have reasonable capture of all major cell classes, including microglia, in our dataset. We then leveraged this dataset to specifically investigate changes in microglial dynamics in the SN from PD and NPC donors, as well as across several regions from PD brains.

### Identification of Human Microglial Subpopulations in PD

We constructed an integrated catalog of CNS myeloid cell responses associated with PD by characterizing the heterogeneity of myeloid cells across our dataset, including cells from all sampled PD regions and the SN of NPCs. As such, we identified all populations expressing the canonical microglial/myeloid markers including *PTPRC*, *CD14, CSF1R*, *AIF1*, and *C1QA* (Fig. 1b and 1c). We then performed unsupervised clustering analysis on this fraction of the dataset to identify distinct subpopulations of microglial/myeloid cells. This process was iterated three times to remove small groups of non-myeloid cells. The final filtered myeloid dataset was then clustered using multiple parameter values to identify distinct clusters (see Methods); the optimal parameter set yielded 15 transcriptionally distinct myeloid subpopulations (Fig. 2a and 2b), ranging in size from 8,767 nuclei to 245 nuclei (Extended Data Fig. 2). We examined the similarity of gene expression between nuclei assigned to different clusters to investigate inter-cluster relatedness and cluster robustness with a post-hoc machine learning approach utilizing a multilayer perception classifier^24, 37, 38^. Using this approach, we found the proportion of ambiguously classified nuclei between clusters ranged between 0% to 6% (Extended Data Fig. 3). As this was below our cutoff value of 20%, we concluded our assignment of cluster identities to be stable with respect to inter-cluster ambiguity at the individual nucleus level.

**Figure 2.**
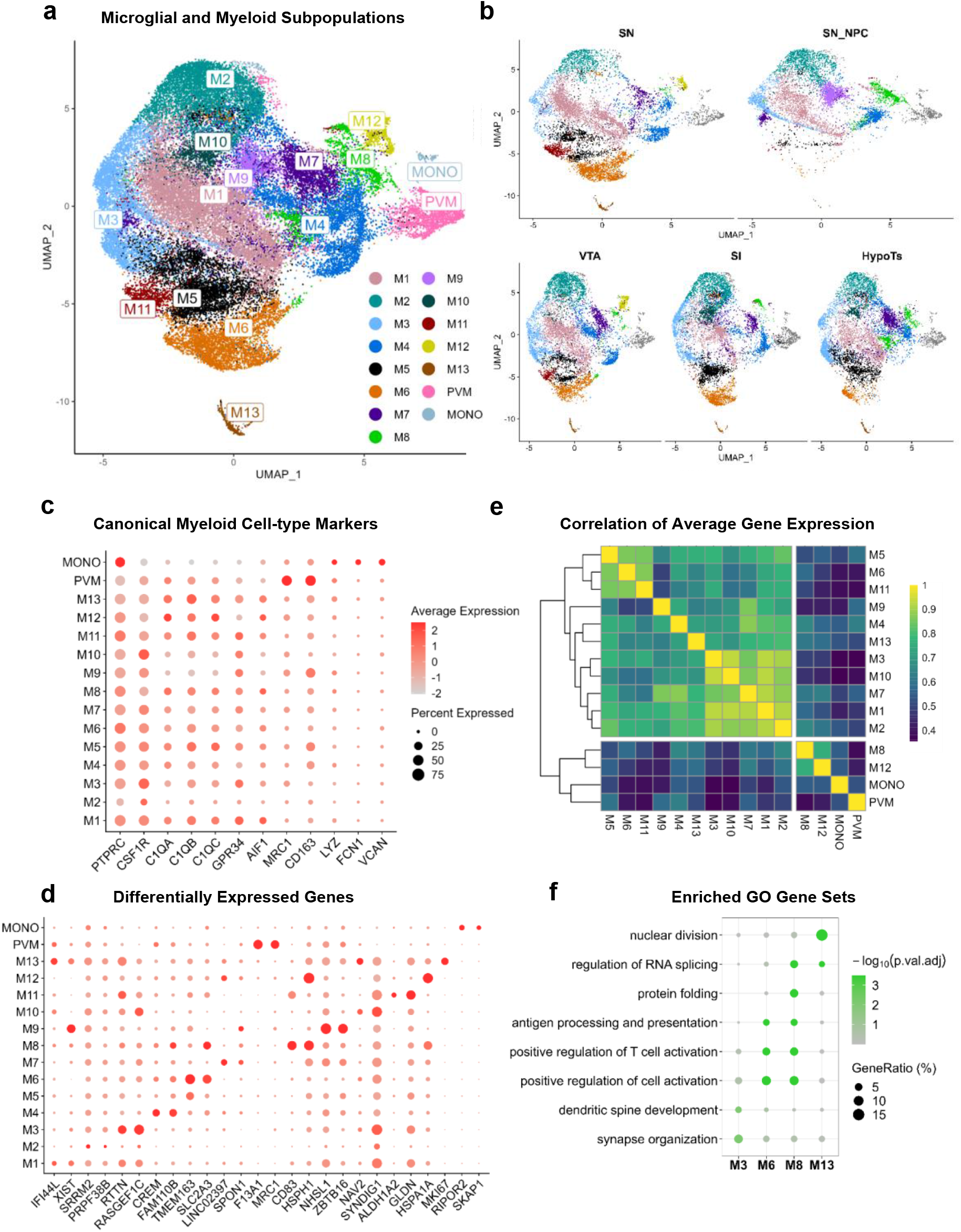
Identification of microglial subpopulations. **a.** Identification of transcriptionally unique subpopulations shown in a UMAP plot of microglial/myeloid nuclei from snRNA-seq data from Parkinson’s disease (PD) and non-PD control (NPC) donors. Clustering analysis using a known nearest neighbor-based approach was used on the identified microglial/myeloid fraction of the total dataset (Fig. 1b). Nuclei from all tissue regions from PD donors including the substantia nigra (SN), ventral tegmental area (VTA), substantia inominata (SI), and hypothalamus (HypoTs), and the SN from NPCs are included in the UMAP plot. Each dot represents a nucleus, and dots are color-coded to indicate subpopulations. **b.** UMAP plot of snRNA-seq data showing the 15 identified microglial/myeloid subpopulations, split by tissue region of origin, including SN, VTA, SI and HypoTs from PD, and SN from NPCs (SN_NPC). **c.** Gene expression pattern of selected general myeloid and canonical microglial, macrophage, and monocyte markers. Each row represents a subpopulation, and each column represents a selected key gene. **d.** Differentially expressed markers for each subpopulation using the Seurat *FindAllMarkers* function. Gene expression of top differentially expressed markers for each subpopulation is depicted. Each row represents a subpopulation and selected genes are represented by each column. **e.** Heatmap of pairwise gene expression correlations (Pearson’s R) of each cluster over the 2,000 most highly variable genes across all the populations. Correlation is colored from low (blue) to high (yellow). **f.** Gene ontology (GO) annotation was performed using the DAVID platform, using biological process annotation. Selected differentially expressed gene-sets are depicted for M3, M6, M8, and M13. GO annotation for M3 reveals homeostatic functional programs, for M6 and M8 enrichment of antigen presentation and T-cell activation gene sets, for M8 unique enrichment for gene sets involved in protein folding and regulation of RNA splicing, and for M13 a proliferative state.

Of the 15 myeloid clusters, all expressed the immune cell marker *PTPRC,* as expected based on the initial subsetting of these nuclei. 13 clusters expressed high levels of microglial genes including *CSF1R*, *C1QA, C1QB, C1QC, GPR34,* and *AIF1* but did not express markers associated with other myeloid cell types (Fig. 2c). We therefore annotated these 13 clusters as subpopulations of microglia which we have labeled as M1-M13, ordered in size from largest to smallest. The two remaining clusters appear to be non-microglial, myeloid populations. One cluster clearly expresses canonical markers of perivascular macrophages, including *MRC1,* as well as the macrophage marker *CD163.* Accordingly, we labeled this population as “PVM”. Additionally, the smallest cluster has minimal *CSF1R* expression, and expresses monocyte-enriched genes including *LYZ*, *FCN1*, and *VCAN*, suggesting that this cluster represents monocytes; accordingly, we labeled it as “MONO”.

After identifying which of our clusters were likely bona fide microglia, we characterized their individual transcriptional signatures. First, we identified genes whose expression was enriched in each microglial subpopulation (see Methods) to determine differences in their functional programs (Fig. 2d; refer to Supplementary Table 3 and Supplementary Table 4 for lists of differentially expressed genes). To determine the similarity between the transcriptional profiles of clusters, we also performed pairwise gene expression correlations (Pearson’s R) of each of the clusters over the 2,000 most highly variable genes across all the populations (Fig. 2e). As M1, M2, and M3 are the most abundant microglial clusters and differentially express markers associated with homeostatic microglia, including *P2RY12* and *CX3CR1,* we posit that they represent homeostatic microglial clusters^39^. Furthermore, clusters M1, M2, and M3 have average gene expression correlation scores of >0.9 for each pairwise correlation among them, suggesting that they are highly transcriptionally similar subpopulations. Clusters M7 and M10 also appear to be homeostatic and transcriptionally similar to M1, M2, and M3, with average gene expression correlation scores of >0.84 to each of these three clusters. Specifically, cluster M3 is strongly enriched for established homeostatic microglial genes *P2RY12* and *CX3CR1.* Indeed, gene ontology (GO) analysis on its differentially expressed markers (Fig. 2f) demonstrates enrichment for gene sets involved in dendritic spine and dendrite development (adjusted p=4.6e-03, adjusted p=7.9e-03) and synapse organization (adjusted p=1.4e-02)^40, 41^. Finally, cluster M2 has lower gene and transcript detection counts, which could represent either a biologically quiescent state or a group of lower-quality nuclei.

Clusters M5, M6, and M11 also have transcriptionally similar signatures, with M5 and M6 sharing average gene expression correlation scores of 0.86, M5 and M11 scoring 0.85, and M6 and M11 scoring 0.87 (Fig. 2e). Interestingly, M5 and M6 are both highly enriched for *TMEM163*, a PD genome-wide association study (GWAS) hit that encodes a zinc efflux transporter, with a log2 fold change (LFC) of 1.2 and 2.4, respectively, compared to all other microglia^42, 43^. Clusters M5 and M6 also have higher expression of genes associated with the inflammatory response and myeloid cell activation, including *SPP1*, *IL17RA, IL1RAP, NAIP*, and *HCK* (Fig. 2f). Despite these overlapping signatures, M5 and M6 do have some notable differences. M6 has higher LFC expression of all the mentioned inflammatory markers compared to M5, and differentially expresses additional inflammatory markers including *IL1B, CASP4, IFI16*, and *TLR2*. Interestingly, 99.7% of nuclei that comprise M6 originate from male donors. Further validation in a larger cohort is warranted to determine whether this subpopulation is sex specific.

M8 strongly differentially expresses genes involved in antigen presentation, the response to misfolded proteins, and RNA splicing, all of which are of particular interest in PD pathogenesis^44–46^. Indeed, this cluster is consistently enriched for heat shock proteins, including *HSPH1, HSP90AA1, HSPB1, HSPD1, HSP90AB1, HSPA1A, HSPA1B, HSPE1, HSPA5, HSPA8,* and *HSPA9,* as well as other genes involved in the response to unfolded proteins, including *DNAJB1* and *BAG3.* Indeed, GO analysis on M8’s differentially expressed genes (Fig. 2f) demonstrates enrichment for genes associated with response to unfolded protein (adjusted p=1.00e-06). M8 is also enriched for genes involved in RNA splicing (GO analysis adjusted p=1.23e-06), including *TRA2B, CLK1, YTHDC1, SRSF3,* and *HNRNPH1* (Fig. 2f). Interestingly, M8 microglia are also highly enriched for *NR4A2* (LFC=1.67) compared to all other microglia, which has been genetically associated with familial PD^47^. Its protein product, NURR1, has also been implicated in the maintenance of dopaminergic neurons^48, 49^.

Furthermore, M8 is enriched for several genes involved in antigen presentation, including HLA genes *HLA-A, HLA-B, HLA-DBQ1, HLA-DPA1, HLA-DBQ1, HLA-DQA1, HLA-DBP1, HLA-DRB1, HLA-DRB5, HLA-DRA*, and others, as well as *CD83*, *CD74, and CTSB*. Several of these genes including *BAG3, CTSB, HLA-DQB1,* and *HLA-DRB1* have been associated with PD through GWAS and fine mapping^50, 51^. Notably, M8 is dramatically enriched for *CD83* (LFC=3.72), which is known to stabilize MHC-II and CD86 expression at the cell surface as well as play an immunoregulatory role^52, 53^. To a lesser degree than M8, M6 also differentially expresses antigen presentation markers including seven of these HLA genes, suggesting that both M8 and M6 are involved in antigen presentation. Importantly, heat shock proteins are essential for antigen presentation, so enrichment in M8 for genes involved in protein folding and antigen presentation may be related to the functional program of this microglial subpopulation. However, as there is a proteinopathy component to PD, enrichment for chaperone proteins may also have importance in this functional capacity.

M9 differentially expresses *GPNMB* (LFC=1.14) and *LRRK2* (LFC=1.17). Interestingly, M9 also differentially expresses *CD163* (LFC = 1.19), as does M5 (LFC=0.94), which is a marker thought to be specific to macrophages and not expressed on microglia^54–57^. It warrants further validation to determine whether these clusters may originate from infiltrating peripheral cells that differentiate into monocyte-derived microglial-like cells once they take up residence in the CNS. M12 appears to be specific to one donor, as 99.5% of nuclei from M12 originate from one individual. Finally, M13 differentially expresses *MKI67* and is enriched for genes involved in cell division (Fig. 2f), suggesting that it may define a pool of proliferating microglial cells.

We investigated each of these microglial subpopulations for enriched expression of PD susceptibility genes using the genes nominated by PD GWAS^50^. We found M3, M6, and M8 to be significantly enriched in PD susceptibility genes (Supplementary Table 5), highlighting a possible role for these subpopulations in PD disease processes.

Thus, the above analysis indicates that our single nucleus dataset offers a robust myeloid cell structure, comprised of microglia, PVMs, and monocytes. The thirteen microglial subpopulations we have identified are likely functionally distinct, as evidenced by their patterns of differentially expressed genes, which fall into three main categories: homeostatic, proinflammatory, and stress-response.

### Unique Microglial Populations are Associated with the SN of PD and NPC Donors

As the SN is a main target of the neurodegenerative process in PD^1^, we leveraged our generated brain myeloid cell structure to investigate whether any clusters were differentially represented in this specific brain region between PD donors and NPCs using mixed-effects association of single cells (MASC)^58^ analysis, controlling for donor, age, and sex as covariates. From this analysis, we found that M2, M8, and MONOs are depleted in the SN of PD donors (Fig. 3a and 3b). Notably, the odds ratio (OR) of nuclei from M8 originating in the SN of PD donors compared to NPCs is 0.125 (95% CI [0.1248, 0.1251], p=0.018), demonstrating a profound depletion of this subpopulation in PD. Since M8 is enriched for transcripts of chaperone proteins and genes involved in the response to unfolded proteins and antigen presentation, its dramatic depletion in PD suggests that a microglial-mediated response against misfolded protein pathologies may be impaired. Conversely, we observed that the proliferative cluster M13 is enriched in the SN of PD donors compared to NPCs. It has been shown that neurodegenerative cues can drive proliferation of selected microglial populations, a process which may underlie the enrichment of proliferating microglia in the SN in PD compared to NPCs^59^.

**Figure 3.**
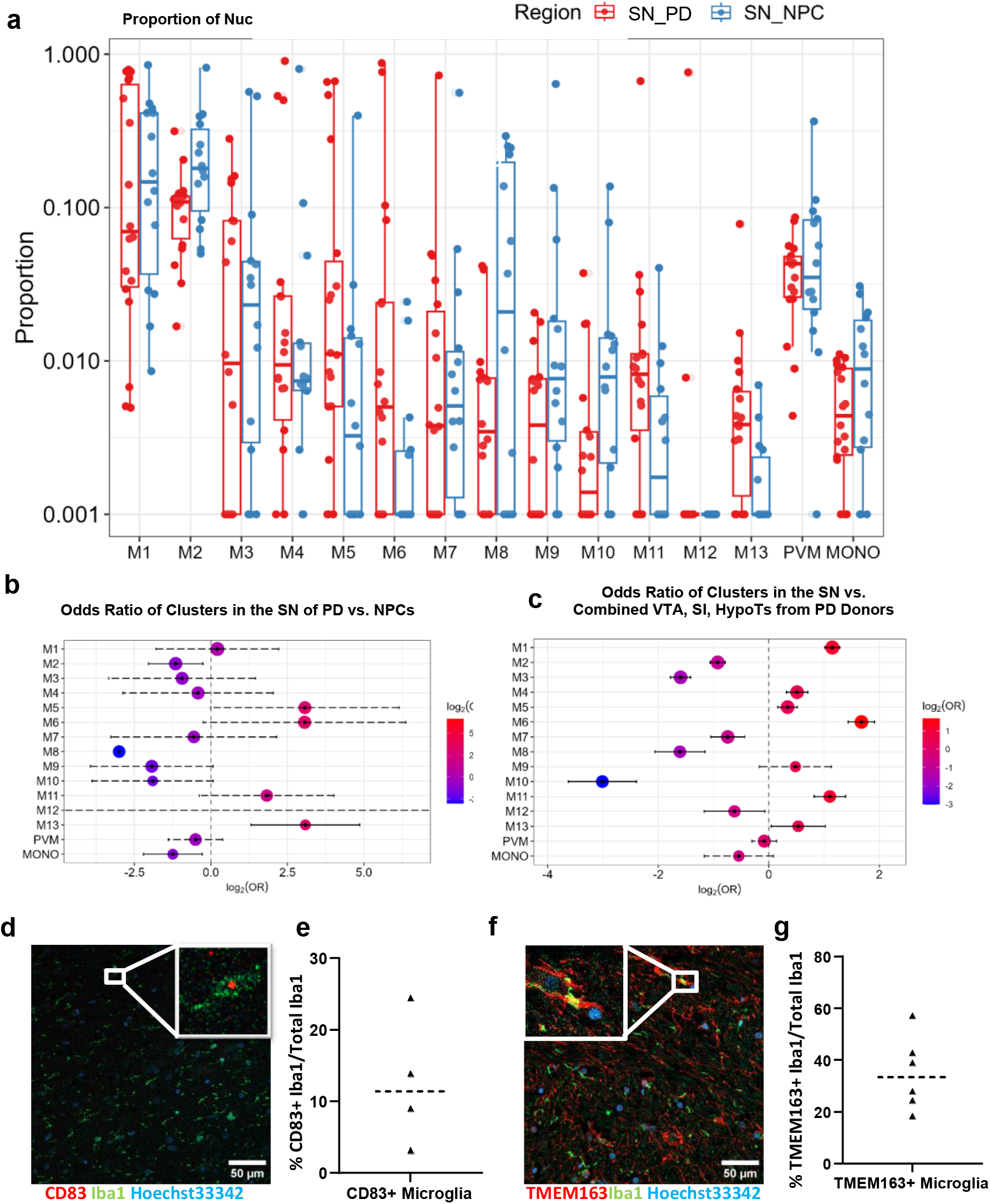
Differential representation of microglial populations across disease conditions and regions. **a.** Box plot of the proportion of nuclei in each cluster from each individual substantia nigra (SN) tissue sample. Nuclei proportions in each cluster are split by disease condition, Parkinson’s disease (SN_PD, red) or non-PD control (SN_NPC, blue). **b.** Proportion of clusters between the SNs of PD and NPC donor samples using mixed-effects association of single cells (MASC) analysis, controlling for donor, age, and sex as covariates. Forrest plot represents the odds ratio of each nucleus within a given cluster originating from the SN of PD vs NPC donors. Color scale indicates enrichment (red) or depletion (blue) of a cluster in the PD SN compared to NPCs. **c.** Proportion of clusters between the SN of PD donors and all PD regions combined (SN, ventral tegmental area [VTA], substantia inominata [SI], and hypothalamus [HypoTs]) using MASC analysis, controlling for donor, age, and sex as covariates. Forrest plot represents the odds ratio of each nucleus within a given cluster originating from the SN of PD vs all other PD regions combined. Color scale indicates enrichment (red) or depletion (blue) of a cluster in the PD SN compared to all other PD regions. **d.** Representative image of *in situ* validation of M8 by immunohistochemistry for CD83+/Iba1+ microglia in the SN of NPCs. Scale bar = 50μm. **e.** Percentages of CD83+ microglia of total Iba1+ microglia for each NPC SN tissue assayed with immunohistochemistry. Dashed line represents the median. **f.** Representative image of *in situ* validation of M5 and M6 with immunohistochemistry of TMEM163+/Iba1+ microglia in the SN of PD donors. Scale bar = 50μm. **g.** Percentages of TMEM163+ microglia of total Iba1+ microglia for PD SN tissue assayed with immunohistochemistry. Dashed line represents the median.

As M6 is nearly entirely comprised of nuclei from male donors, we further investigated whether there were any differentially represented microglial populations in the SN of PD versus NPCs in the male donors in the cohort. Using MASC analysis and controlling for age and donor as covariates, we found M2, M8, and MONOs are similarly depleted within the male donor subset, and that M6 and M13 are enriched in the SN of PD donors (Extended Data Fig. 4a and Extended Data Fig. 4b). Indeed, the OR of nuclei from M6 originating from the SN of PD donors compared to NPCs is 16.6 (95% CI [1.38, 199.33], p=0.03), signifying that this microglial population is specific to the PD SN in these male subjects. Thus, the above results reveal divergences between microglia in the SN of PD donors versus NPCs. Some subpopulations of microglia, particularly M8, are depleted in the SN of PD donors compared to NPCs. Others are more prevalent in PD, one of which, M6, may be sex-specific.

Next, we elected to validate these clusters *in situ* by immunohistochemistry on tissue sections from the PD and NPC subjects used for single-nucleus sequencing (see Methods). We probed SN tissues from four NPCs for CD83+ microglia, a marker that is highly specific to M8 at the RNA level within our dataset, with a receiver operator characteristic (ROC) classification power score of 0.89. From the immunohistochemistry of NPC SN tissues, we observed CD83+ microglia at a frequency of 3-24% (mean = 12.7%) of all Iba1+ microglia (Fig. 3d and 3e). This corresponds to the frequency of M8 observed in these same four individuals from our dissociated nuclei, ranging from 4-28% (mean=19.5%) of microglia. We also probed SN tissues from six male PD donors for TMEM163+ microglia, the top enriched marker for M5 and M6. We observed TMEM163+ microglia in the SN from these six individuals at a frequency of 18-57% (mean=35%) of all Iba1+ microglia (Fig. 3f and 3g). This proportion is within the frequency range of combined M5+M6 nuclei observed in the SN from these same individuals of 6-87% (mean=53.0%). Thus, these immunohistochemical data confirm the presence of these microglial subpopulations in the SN of PD donors and NPCs.

### Regional Specificity in PD Microglial Subpopulation Dynamics

Next, in addition to examining differences in microglial subpopulations between SN tissues from PD donors and NPCs, we also investigated whether there were any differences in cluster representation across brain regions in PD with different degrees of neuronal loss. We thus performed MASC^58^ analysis on nuclei from the SN and the three other brain regions selected from PD cases (VTA, SI, HypoTs; Supplementary Table 2 and Extended Data Fig. 1), controlling for sex, age, and donor. We found that compared to nuclei from all other regions combined (VTA, SI, and HypoTs), the SN of PD donors has lower proportions of M2, M3, M7, M8, and M10, and higher proportions of M1, M4, M5, M6, M11, and M13 (Fig. 3C). In particular, the lower prevalence of chaperone enriched M8 (OR =0.33, 95% CI [0.24, 0.45], p<0.001), mirrors that in the SN of PD donors compared to NPCs. This finding confirms regional depletion of NPC-associated microglial programs in a manner topographically associated with disease. Interestingly, we also observed focal enrichment in the SN of PD donors for the *TMEM163* high clusters M5 and M6 as well as M13 compared to other PD regions, mirroring the pattern observed in comparison to NPCs. This regional analysis demonstrates that compared to all other PD tissues (combined VTA, SI, and HypoTs) the SN in PD donors shows strong differences in microglial subpopulations that are reminiscent of those observed when compared to NPCs – suggesting that our findings represent microglial dynamics that are strongly associated with PD neurodegeneration.

To further investigate regional variation of the M8 subpopulation depletion in PD, we performed pairwise MASC analysis between each PD region, controlling for age, sex, and donor. We consistently found that M8 was increasingly reduced in the SN compared to the VTA (Extended Data Fig. 5a; OR=0.35, 95% CI [0.26, 0.49], p<0.001), the SI (Extended Data Fig. 5b; OR=0.1, 95% CI [0.072, 0.143], p<0.001), and the HypoTs (Extended Data Fig. 5c; OR=0.05, 95% CI [0.036, 0.069], p<0.001). Thus, the relative abundance of M8 appears to be associated with the relative vulnerability of each region to PD neurodegeneration. This finding further suggests that the regional reduction in M8 microglia parallels the magnitude of the neurodegenerative process.

### Validation of M8 Within a Non-Diseased Human Brain Transcriptomic Dataset

Considering the striking depletion of M8 between the SN in PD and NPC samples, as well as across our selected brain regions in PD, we sought to investigate the regional dynamics of M8 in non-diseased adult brains using the dataset from Siletti et al. (2022), who generated snRNA-seq data from 10 different brain regions of three individuals without neurological disease^36^. We integrated all nuclei from the SN of NPCs from our myeloid dataset (15 clusters) with all microglial nuclei from the Siletti et al. dataset (Fig. 4a). Visually, we observed the nuclei from the SN of NPCs from our dataset overlapped well with nuclei from the Siletti dataset in the uniform manifold approximation and projection (UMAP) space. We then performed clustering analysis using this integrated dataset (Fig. 4b). Within the integrated dataset, we observed that the majority of M8 nuclei localized to cluster 7, a cluster that is distinct within the UMAP space (Fig. 4c). We validated that this integrated cluster 7 carries the M8 transcriptional signature, finding that the top differential markers for M8 are robustly expressed only within integrated cluster 7 in the combined dataset (Fig. 4d). These findings demonstrate the presence of microglial nuclei carrying the M8 signature within the non-disease brains sampled in the Siletti et al. dataset.

**Figure 4.**
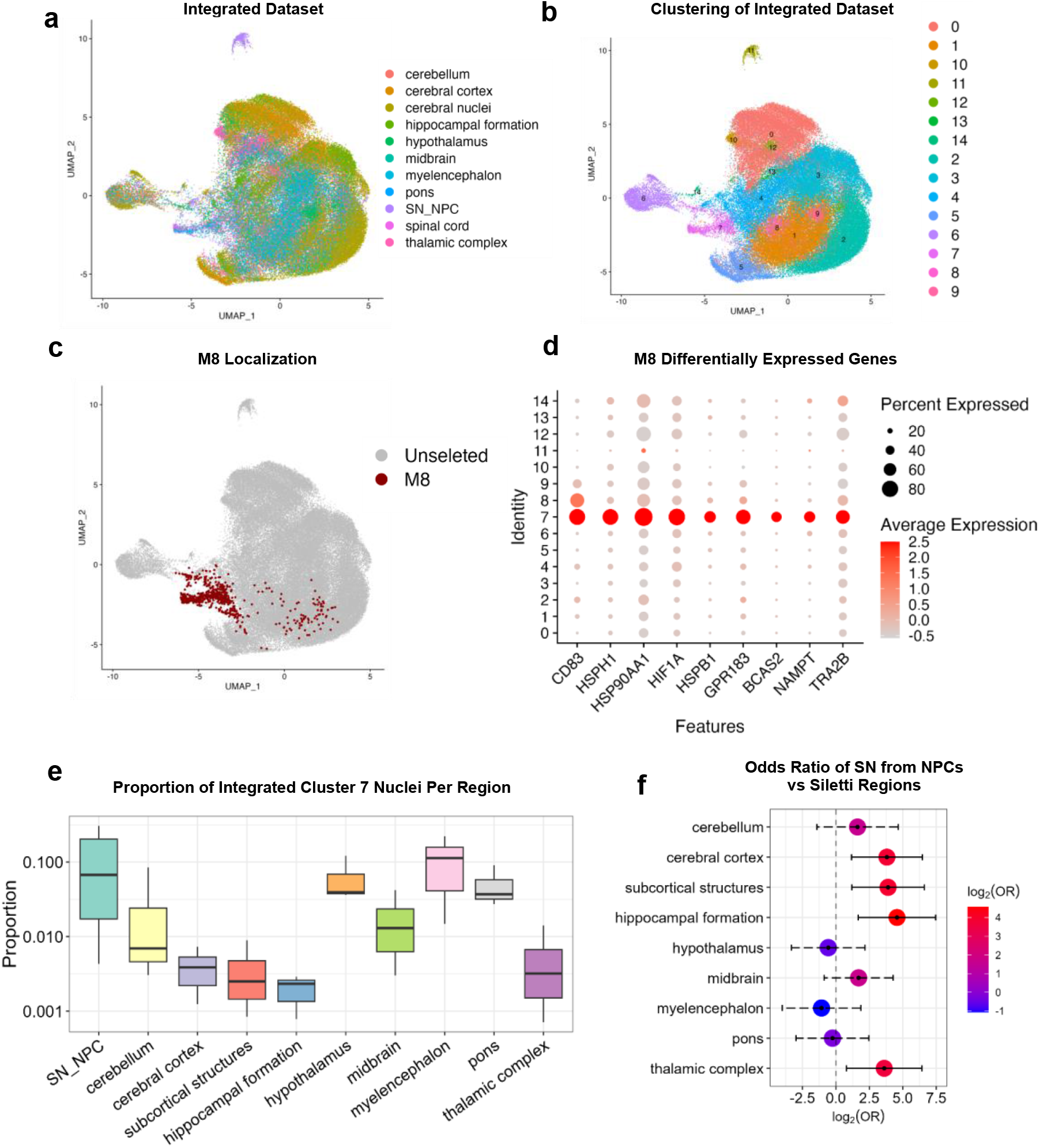
Validation of M8 as a regionally-specific microglial subpopulation in non-diseased human brain tissue. **a.** UMAP plot of the integrated dataset including all nuclei from the SN of non-Parkinson disease controls (NDCs) and all microglial nuclei from the Siletti et al. (2022) dataset^36^, which includes microglia across 10 brain regions from individuals without neurological disease. Each dot represents a nucleus, and dots are color-coded to indicate brain region of origin. **b.** Clustering analysis of the integrated SN_NPC and Siletti et al. dataset using a known nearest neighbor-based clustering approach. The resulting clusters are shown in a UMAP plot, in which each dot represents one nucleus, and each cluster is represented in a different color. **c.** Nuclei originating from M8 from the SN of NPCs are depicted by red dots within the integrated dataset UMAP space, with all other nuclei depicted as grey dots. M8 nuclei primarily localize with the integrated cluster 7 (Fig. 4b). **d.** Gene expression of the top M8 differentially expressed markers by each cluster in the integrated dataset. Each integrated cluster is represented by a row, with each selected gene represented by a column. The integrated cluster 7 alone robustly expresses all M8 marker genes. **e.** Box plot representing the proportion of cluster 7 nuclei from the integrated dataset within each brain region, normalized across the number of donors contributing nuclei to each of these regions. **f.** Pairwise mixed-effects association of single cells (MASC) analysis comparing the proportion of integrated cluster 7 in the SN of NPCs to every other region in the Siletti et al. dataset. Odds ratios are depicted in a forest plot, with each row representing the pairwise comparison of cluster 7 nuclei in the SN of NPCs compared to the indicated region from the Siletti dataset. Color scale indicates enrichment (red) or depletion (blue) of cluster 7 in the SN of NPCs compared to each of the indicated regions.

To investigate regional differences in the M8 signature, we calculated the proportion of integrated cluster 7 nuclei (as a fraction of the total microglia) within each brain region, per donor in the Siletti et al. dataset (Fig. 4e). We performed pairwise MASC analysis to investigate regional differences in cluster 7 distribution (Fig. 4f). We found that the median proportion of cluster 7 nuclei is higher among tissues originating in caudal brain regions, including our NPC SN samples, as well as midbrain, pons, medulla oblongatta (myelencephalon), hypothalamus, and cerebellum samples from the Siletti et al. dataset (Fig. 4e). MASC analysis demonstrated that integrated cluster 7 nuclei carrying the M8 signature were enriched in nuclei from the SN of NPCs (Fig. 4f) compared to cortical (OR =14.08, 95% CI [2.24, 88.32], p=0.02), subcortical (OR = 14.95, 95% CI[2.29, 97.30], p=0.03), hippocampal (OR = 23.77, 95% CI [3.19, 176.77], p=0.01), and thalamic regions (OR=12.27, 95% CI [1.74, 86.61], p=0.04). No difference between the proportion of cluster 7 nuclei was found between the SN of NPCs compared to the midbrain, pons, medulla oblongatta, hypothalamus, or cerebellum. These findings suggest that microglia carrying the M8 signature may have regional specificity to the hypothalamus, cerebellum, and brainstem, which notably includes the most affected regions in PD.

### M8 Has Unique Open Chromatin Compared to Other Microglial Subpopulations

We next sought to integrate the ATAC- and RNA-seq signatures in order to identify cluster-specific chromatin configurations within our described microglial subpopulations, as well as identify putative transcription factors involved in their transcriptional regulation. Accordingly, we assessed chromatin signatures through the ATAC-seq data on these same nuclei in our microglial/myeloid dataset. A total of 29,122 nuclei were considered within the ATAC analysis (see Methods for quality control, Extended Data Fig. 6).

We clustered the ATAC-seq data using the ArchR framework, which also incorporates the Seurat *FindCluster* function (Fig. 5a). In this way, we identified 8 unique ATAC-seq clusters within our microglial/myeloid dataset. When we assigned the RNA-seq cluster labels to this analysis, we observed that four RNA-seq clusters also appear to cluster separately within the ATAC space: PVMs, MONOs, M8, and M11 (Fig. 5a and Extended Data Fig. 7). While it is expected that PVMs and monocytes will have unique chromatin signatures, strikingly, M8 appears to be unique from other microglia in its chromatin structure.

**Figure 5.**
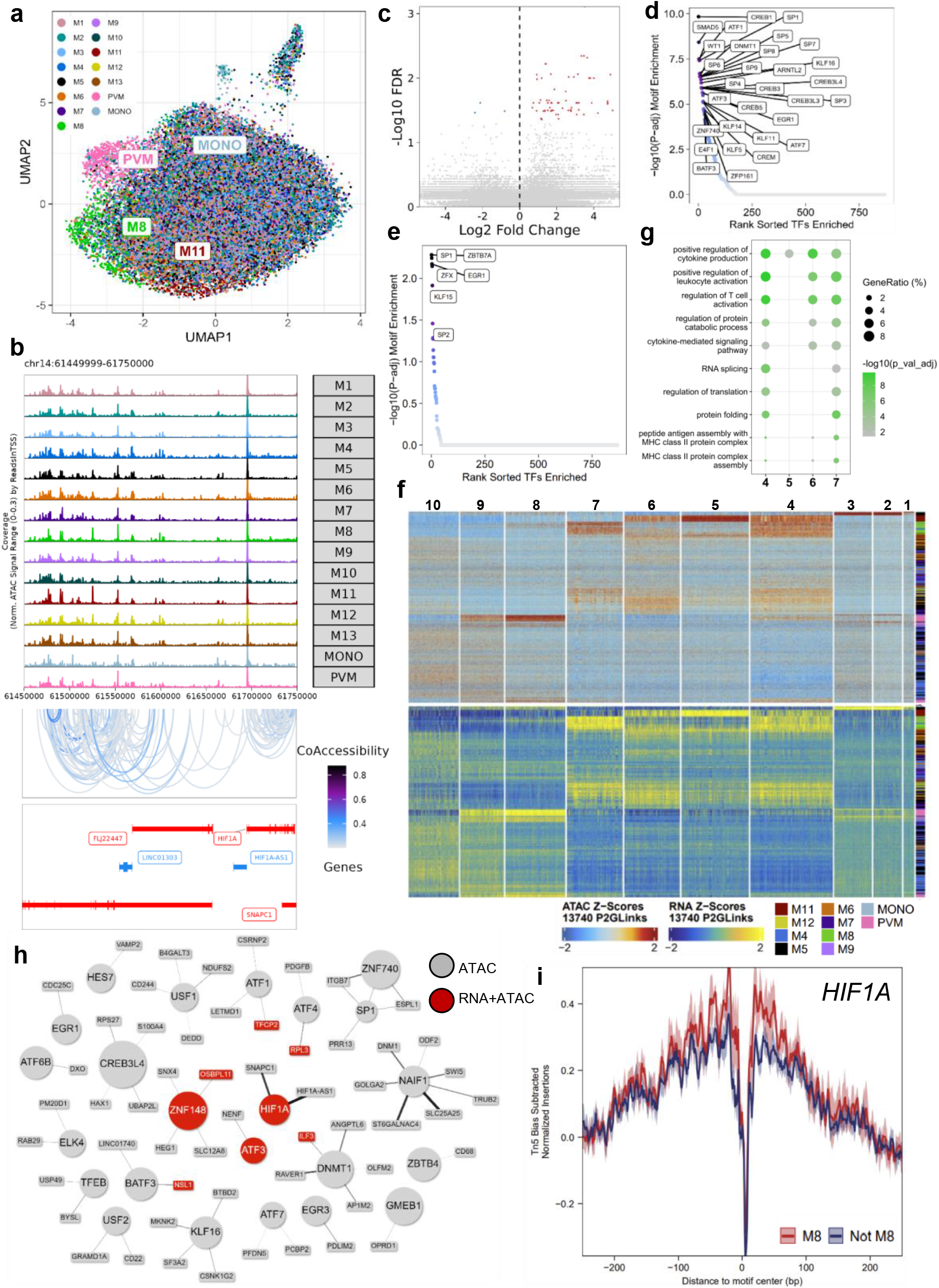
M8 has a unique open chromatin structure compared to other microglial subpopulations. **a.** Clustering analysis of snATAC-seq data performed within the ArchR framework, depicted in UMAP representation. Each nucleus is represented by a dot and annotated per its snRNA-seq cluster label, depicted by color code. Nuclei from all tissue regions from Parkinson’s disease (PD) donors including the substantia nigra (SN), ventral tegmental area (VTA), substantia inominata (SI), and hypothalamus (HypoTs), and the SN from Non-PD controls (NPCs) are included in the UMAP plot. **b.** *<u>Top panel</u>*: depicts accessibility browser track in the neighboring region of the *HIF1A* gene. *<u>Middle panel</u>*: depicts peak co-accessibility within the region with a minimum correlation of 0.2. *<u>Bottom panel</u>*: depicts the known location of genes in the genomic track within this region. **c.** Differentially accessible peaks were identified for each microglial subpopulation. M8 is found to have 63 differential peaks, as shown in a volcano plot of differential accessibly peaks for M8 as compared to remaining clusters with a maximum adjusted p-value of 0.05. **d., e**. Scatter plots highlighting the results of motif enrichment analyses for both up-regulated (**d**) and down-regulated (**e**) accessible peaks for M8 compared to remaining clusters. **f.** Heatmap showcasing 13,740 peak-to-gene (P2G) links, which are correlations between peak accessibility and gene expression. Post-hoc known nearest neighbor clustering analysis of P2Gs as expressed in both RNA-seq and ATAC-seq modalities. Homeostatic clusters are excluded from this heatmap to allow a more refined visualization of the remaining microglial clusters. **g.** Gene ontology analysis for P2G link module 7 (depicted in the P2G heatmap from Fig. 5F). **h.** Network plot showing the highest correlated P2G links for M8, with genes that share the same peak connected to one another. In these connections, chromatin accessibility of enriched motifs from Fig. 5d & 5e was correlated with gene expression resulting in P2G links. Motifs are shaped as circular nodes, while linked genes are shaped as rectangular nodes. Genes that are differential to M8 in the RNA-seq data are colored red. Edge thickness indicates a higher number of P2G links. **i.** Footprint plots for the *HIF1A* motif, which is enriched within the differentially upregulated peaks for M8. M8 footprinting is depicted in red, and footprinting of all other microglial subpopulations is depicted in blue.

Within the ATAC-seq dataset, we observed 240,000 peaks of accessible chromatin (Fig. 5b). We identified differentially upregulated peaks within M8, indicating open accessibility of DNA, using a null hypothesis rejection cutoff threshold of 0.05 after p-value adjustment. In M8, we observed 63 differentially up-regulated (accessible) peaks and 3 down-regulated (less accessible) peaks (Fig. 5c and Supplementary Table 6). We next sought to identify unique transcription factors which may bind within M8-enriched accessible regions. We found 60 motifs enriched within the differentially upregulated peaks for M8, including *HIF1A, NAIF1,* and *BATF3* (p<0.05; Fig. 5d and 5e).

Next, we investigated the relationship between gene expression and chromatin accessibility using the combined multiome ATAC and RNA modalities, to better understand which chromatin signatures may be actively influencing the transcriptional landscape of each microglial subpopulation. We identified accessible chromatin peaks that were highly correlated with gene expression, or peak-to-gene (P2G) links. We sorted these P2G links into 10 modules representing unique patterns of correlated RNA gene expression/chromatin peaks (Fig. 5f). We then leveraged these P2G correlations to investigate which of our identified snRNA-seq subpopulations share unique patterns across both RNA and ATAC modalities. RNA-seq clusters that appear to maintain cohesion in these highly correlated patterns between RNA-seq and ATAC-seq signatures include PVMs, M8, and M11. The chromatin signatures of these three subpopulations are preserved at the transcriptional level and may be driving their individual transcriptional signatures.

Within the 10 identified P2G modules, module 7 contains P2G links that are highly expressed in M8 and appear to be unique to this subpopulation (Fig. 5f). To investigate the functional implications of the correlated high chromatin accessibility and gene expression pattern of P2G-module 7, we performed GO analysis on the genes that had high P2G correlations (Fig. 5g). Interestingly, both the P2G-module 7 GO analysis and the RNA-seq GO analysis for M8 highlighted shared enrichment for genes involved in protein folding, antigen presentation, RNA splicing, and the regulation of T cell activation (Fig. 2f and Fig. 5g).

To better understand the chromatin/transcriptional network influencing the M8 signature, we leveraged the P2G links with the highest correlations within M8 to create a network of transcription factors and other linked genes corresponding to accessible peaks (Fig. 5h). We also investigated which highly correlated P2G links for M8 corresponded to differentially expressed genes (from our RNA-based AUROC analysis) within the snRNA-seq data. From this network, *HIF1A*, *ZNF148,* and *ATF3* are highly correlated P2G links within M8 which are also differentially expressed at the RNA level with high specificity to M8. Of these transcription factors, only *HIF1A* is also found within a differentially accessible peak for M8, suggesting that this motif is also specific to this subpopulation at the chromatin level. We performed transcription factor footprinting analysis for *HIF1A* to investigate transcription factor binding patterns between M8 and other microglial clusters (Fig. 5i). M8 shows elevated levels of binding footprints around the *HIF1A* motif compared to other clusters, suggesting a regulatory role of *HIF1A* that is unique to M8. Given that HIF1α facilitates myeloid cell-mediated inflammation and metabolic reprogramming, this suggests a unique activation state for M8^60, 61^. These findings identify *HIF1A,* amongst other transcription factors and linked genes within the accessible chromatin peaks of M8, as a potential drivers of the unique M8 transcriptional signature.

## Discussion

To gain insights into the role of microglia in PD, we used snRNA-seq and ATAC-seq to generate a multiomic microglial signature that we leveraged to elucidate microglial heterogeneity in this disease. We created this dataset using tissues from the SN of PD donors and NPCs, as well as from the VTA, SI and HypoTs of PD donors to probe regional patterns of microglial dynamics in PD. We identified 13 microglial subpopulations within these tissues with unique molecular signatures and established their relative abundance (M6, M13) or paucity (M8) in the SN in PD. We found that the proinflammatory microglial subpopulation M6 is more prevalent in the SN from PD donors compared to that from NPCs. In PD samples, M5 and M6 are enriched in the SN compared to the VTA, SI, and HypoTs. Conversely, we found that the *CD83* and *HIF1A*-enriched stress-response microglial subpopulation, M8, is diminished in the SN from PD donors compared to the SN of NPCs. Furthermore, M8 paucity appeared to parallel the magnitude of neurodegeneration across our four selected PD brain regions. The reduction of this population may have implications for neuronal vulnerability in disease.

Our identified proinflammatory microglial subpopulations (M5, M6, M11) likely correspond to the reactive microglia observed in the SN of PD patients, which have been reported to express adhesion and inflammatory markers including MHC-II and TLR2^62–65^. These markers are differentially expressed at the RNA level by M6 and the related M11 microglia in our dataset. Moreover, M5 and M6 microglia have enriched expression of the PD GWAS hit *TMEM163,* which encodes a zinc efflux transporter protein and provides a novel specific marker for these subpopulations^42, 43, 66, 67^. *TMEM163* has also been associated with PD as a rare variant, as well as through transcriptome-wide association studies^43, 68, 69^. This finding raises the question of how zinc efflux may impact microglial-mediated inflammation in the PD-SN, as well as how microglia functionally contribute to the PD risk associated with *TMEM163*. Interestingly, neuronal zinc release has been shown to increase under many pathological conditions^70–72^. Furthermore, zinc has been shown to induce microglial NF-κB activity and alter microglial cytokine production, in part through P2X receptor activity, suggesting one possible mechanism through which TMEM163 activity may contribute to the M5/M6 transcriptional phenotype we have observed^71, 73–75^. Additionally, *TMEM163* itself has specifically been demonstrated to regulate P2X7 receptor activity, which has implications for microglial responses to ATP as well as inflammasome activation^76^. We additionally detected enrichment for gene sets in M5 and M6 microglia involved in the inflammatory response and myeloid cell activation, which can be validated at the protein level to better characterize the participation of these subpopulations in neuroinflammation and disease.

In aggregate, these findings raise the question whether the emergence of this proinflammatory microglial phenotype preferentially in the SN of PD donors develops merely in response to the more severe neurodegeneration taking place in this region, or rather is due to a particular inter-cellular communication contributing to disease pathogenesis. One possible explanation for the enrichment of M5 and M6 in the PD SN is an expansion of these populations by a proliferative pool in disease. It has been shown that neurodegenerative cues can guide a rapid expansion of only selected microglial clones^59^. Relevant to this idea, we detected a proliferative microglial subpopulation, M13, enriched in the SN from PD donors compared to NPCs. However, whether these proliferative M13 microglia may later adopt an M5/M6 signature remains to be demonstrated. Additionally, nearly all M6 nuclei in our dataset originated from male donors. As males develop PD at a ratio of 3:2 compared to females^77^, replication and further investigation of this finding in a larger cohort may shed light as to whether M6 microglia may contribute to sex differences in PD.

In addition to observing an enrichment in proinflammatory microglia, we conversely observed a stark depletion of M8 microglia in the SN from PD donors compared to that from NPCs. The abundance of M8 microglia in NPC SN tissues prompted us to investigate M8 subpopulation dynamics in the non-diseased brain. Using the Siletti et al. (2022) single nucleus dataset of human brain tissues from donors without neurological disease^36^, we found that nuclei carrying the M8 signature have regional specificity to brain regions including the midbrain, pons, medulla oblongatta, hypothalamus, and cerebellum. This analysis importantly revealed that M8 microglia are enriched in the brainstem, which according to Braak’s staging^3, 30^, is a key and initial locus of PD neuropathology, as well as validated our own observation of M8 microglia within the midbrain of NPCs.

We next investigated whether the magnitude of M8 depletion parallels the extent of damage among our selected PD brain regions. Remarkably, we found that M8 depletion becomes more pronounced with increasing disease involvement, such that M8 microglia are abundant in the SN of NPCs and depleted in the SN of PD donors, but are present in the HypoTs from PD regions. This pattern of depletion is suggestive of M8 loss with neurodegenerative processes. Human and mouse studies have described epigenetic and transcriptional microglial signatures dependent on the environment, and it is possible that M8 microglia form as a result of caudally-specific environmental cues that are lost as a result of PD processes^78, 79^. Indeed, environmental cues from neighboring glia, neurons, and infiltrating molecules from the blood-brain barrier have been shown to contribute to regional heterogeneity in microglia^80^. However, the mechanism of M8 microglial depletion in PD is unclear; whether these cells die or are polarized away from this fate to another microglial signature.

Our findings of M8 depletion in the SN from PD donors raises the possibility that this specific microglial subpopulation may contribute to maintaining the wellbeing of neighboring neurons. One mechanism through which M8 may exert a neuroprotective role is through NURR1 signaling. M8 differentially expresses *NR4A2* (encoding NURR1), which is required for the maintenance of dopaminergic neurons and is genetically associated with familial PD^47–49^. Microglial NURR1 expression and signaling has been shown to inhibit the expression of proinflammatory genes and result in the protection of degenerating dopaminergic neurons through paracrine signaling^81, 82^. Moreover, M8 microglia are potently enriched in genes that have been shown to be involved in the stress-response, antigen presentation, and RNA splicing. M8 microglia are enriched for numerous genes that encode heat-shock proteins (HSPs), including members of the HSP70 family. HSPs are required for antigen presentation, so this may hold implications for the productive handling of proteins in the immune synapse. Thus, this cluster may represent a population of microglia that is responding to stressful stimuli, through antigen presentation and managing folding of proteins, all of which may be instrumental in maintaining a healthy neuronal environment.

Relevant to a unique and potentially neuroprotective functional role for M8 microglia, chromatin signatures of M8 highlight *HIF1A* as an enriched motif, a transcription factor that is also differentially expressed by M8 nuclei at the RNA level. HIF1α is essential for myeloid cell-mediated inflammation, and its enrichment in M8 microglia is suggestive of a unique metabolic state^60, 61^. Enrichment for MHCII genes as well as *SPP1* and *CD74* further suggests an activated role for M8 microglia. Importantly, inflammation is a necessary process for the removal of CNS stimuli that cause neuronal stress as well as repair and recovery mechanisms. Microglia can be activated by misfolded proteins such as α-synuclein in PD and related disorders, and M8 appears to be particularly poised to respond to such stimuli. M8 microglia may provide a functional activated role within the SN in steady-state contexts.

In this work, we have demonstrated the focal depletion of M8 microglia in the SN from PD donors; a subpopulation with molecular signatures that are suggestive of protective functions in PD. An essential question in the field is how to sustain and promote neuroprotective microglial phenotypes as a therapeutic approach. Equally important is developing methods to selectively reduce inflammatory microglial populations without compromising the function of beneficial subpopulations. In this vein, future work integrating our microglial genomic data with neuronal and other glial data to generate a multimodal signature of neuroimmune interactions will help identify putative pathways involved in differential susceptibility of neurons in PD. This work can ultimately be used to develop specific biomarkers and candidate molecules for potential therapies aimed at targeting specific neuroimmune interactions, whether enhancing neuroprotective microglial effects or suppressing microglial signaling that may contribute to neuronal loss, rather than broadly modulating immune function.

## Methods

### Selection of PD and Non-Disease Control Donors and Acquisition of Brain Tissue

Tissue samples from pathologically confirmed PD brains (n = 19) and age and post-mortem delay-matched controls without neurological or neuropathological signs of PD (referred to henceforth as non-PD controls or NPC; n=14) were obtained from the New York Brain Bank at Columbia University Irving Medical Center and were accrued and processed per its published protocol^83^. All PD donors held a clinical diagnosis of PD that was confirmed in autopsy, were levodopa-responsive, and had no other neurodegenerative conditions. PD cases were staged using the regional location of Lewy pathology and had Braak Lewy body stages of IV–VI^3^. Controls were restricted to samples with no antemortem diagnosis of PD, no more than minimal neuronal loss in the substantia nigra, an absence of Lewy bodies, and a Braak neurofibrillary tangle stage ≤IV. Upon removal, brains were transected sagittally: half of the brains were immersed in formalin for neuropathological assessment and diagnosis while the other half were further dissected into 18 standardized brain blocks (SBBs) prior to being flash frozen and stored at −80°C until use. For the PD cases, we selected SBB#7 which contains the substantia inominata (SI) and the rostral hypothalamus (HypoTs; also known as the preoptic area) and SBB#10 which contains the substantia nigra pars compacta (SN) and the ventral tegmental area (VTA). These different regions were selected to provide a range of neurodegeneration which we supported by semi-quantitative assessments of neuronal loss (Extended Data Fig. 1)^30–35^. For the NPCs, SN tissues were also collected. Neuronal loss (NL) was staged from a scale of 0-4, with 0 representing no neuronal loss, 1 representing mild, 2 representing moderate, 3 representing severe, and 4 representing very severe neuronal loss. Lewy bodies were counted on sections from the described regions following immunohistochemistry for alpha-synuclein (Leica, cat# ncl-l-asyn, 1:100-400 dilution). Of note, the neuropathological evaluation of SN was performed at two levels, caudally at the decussation of the superior cerebellar peduncle and rostrally at the level of the red nucleus, and collapsed into one measure. See Table 1 and Supplementary Table 1 and Supplementary Table 2 for details about the demographics and neuropathology of PD and NPC cases. This research protocol was approved by the Institutional Review Board (IRB) of Columbia University Irving Medical Center (protocols # AAAU7125, #AAAB0192).

### Single Nucleus Isolation

Nuclei were isolated from fresh frozen brain tissues. 30-50mg of tissue were used for each sequencing sample. All steps were performed on ice. Tissues were placed in 2mL of Nuclei EZ lysis buffer (Sigma, cat#NUC101) and dissociated manually in two steps using a blade, followed by a dounce homogenizer (Sigma, cat#d8938). Tissues were dounced 20 times with Pestle A and then 25 times with Pestle B, and then transferred to a 15mL conical tube containing an additional 2mL of EZ lysis buffer. Preparations were then incubated in Nuclei EZ lysis buffer on ice for five minutes and centrifuged at 500g at 4°C for 5 minutes to isolate nuclei and remove cytoplasmic contaminants. The resulting pellets were resuspended in nuclei EZ lysis buffer, and the preparations were again incubated on ice for 5 minutes and centrifuged at 500g for 5 minutes. Preparations were then rinsed in a nuclei suspension buffer (NSB; made of PBS, 0.01%BSA and 0.1% RNAse inhibitor (Takara, cat# 2313A)), centrifuged at 500g for 5 minutes, and resuspended in NSB. Preparations were then filtered through a 35uM mesh cell-strainer (Corning, cat#352235) twice to further remove debris.

### Single Nucleus RNA- and ATAC-Seq

Sample batches of nuclei preparations for snRNA-seq and ATAC-seq were structured to contain tissues from different donors across tissue region, as well as disease. Nuclei preparations were sequenced using the 10x Genomics Chromium platform. snRNA/ATAC-seq libraries were prepared according to the 10x Genomics protocol using the Single Cell Multiome ATAC + Gene Expression kit, followed by sequencing on the Illumina Novaseq platform to a targeted depth of 50k reads/nucleus for RNA-seq and 25k reads/nucleus for ATAC-seq libraries. Raw sequencing reads were aligned and quantified to the human GRCh38 reference, with Ensembl transcript annotation GRCh38.91, using the Cellranger ARC v2 software package from 10x Genomics. Subsequently, the CellBender software package (https://github.com/broadinstitute/CellBender) was used to remove putative background/ambient RNA in each sample, with the following parameters: expected-cells=5000, total-droplets-included=20000, fpr=0.01, epochs=50. Downstream QC and clustering analyses (described below) then proceeded using the Cellbender-corrected hdf5 expression files as input.

### Single Nucleus RNA-Seq Quality Control

Downstream analysis was performed using the R programming language and the RStudio environment. 10x Genomics Chromium runs for every human tissue sample underwent an initial filtering process, which employed a cutoff of ≥1,000 nUMIs. In addition, nuclei with more than 10% of UMIs mapping to mitochondrial genes were removed. Before downstream clustering, all mitochondrial genes were then additionally removed from the count matrices as well. After these quality control steps, 481,703 total nuclei were remaining, with a median detection of 2,058 genes per nucleus.

### Clustering and Top-Level Annotation

The snRNA-seq datasets from all 85 tissue samples were merged and then log-normalized using the Seurat v4.0 R package^84^. The merged datasets were then analyzed jointly to determine the optimal Principal Component values, as determined using the *ElbowPlot* and *PCheatmaps* functions. A PC value of 15 was selected for clustering and UMAP visualization. Clusters were identified using a resolution of 0.5, and then annotated manually based on the expression of canonical marker genes, as well as the expression of cell-type specific markers within the differentially expressed genes for each cluster found with Seurat’s *FindAllMarkers* function. Clusters were broadly annotated based on expression of the following markers: neurons (*SNAP25, SLC17A7, SLC17A6, SYT1, TH, GAD1, GAD2*), astrocytes (*AQP4*), microglia (*AIF1, P2RY12, CSF1R, C1QA, PTPRC, CD14*), T-cells (*CD3E, ITK*), oligodendrocytes (*MOBP, MBP, OPALIN*) oligodendrocyte precursor cells (*OLIG1, PDGFRA*), endothelial cells (*FLT1, CD13*) and pericytes (*PDGFRB*). Some populations which did not show a clearly distinct expression pattern of cell type-specific markers were marked “unassigned”.

### Iterative Removal of Mixed Signature Nuclei from Myeloid Populations

After identifying microglia from the top-level annotation, these nuclei were subsetted to form a new microglial/myeloid dataset, on which all downstream analyses were performed. Subclustering analysis was performed with the same workflow used to cluster the larger dataset, as described above. Subpopulations consisting of non-myeloid contaminating cells were identified with the canonical cell-type specific markers detailed above. Following initial subclustering analysis, nuclei with detection of (non-microglial) glial and neuronal transcripts were identified in this way and removed from the dataset. This process was iterated three times until all non-myeloid contamination was removed from the dataset.

### Subclustering of Myeloid Cells

After the iterative cleanup described above, we subclustered the myeloid cells using 15 PCs and default parameters for the *FindNeighbors* function in Seurat. We tested microglia subclustering across values of the Seurat resolution parameter, ranging from 0.2 to 0.5. We selected a cluster resolution of 0.3, as it identified subpopulations with unique combinations of differentially expressed genes distinguishing each cluster.

We next evaluated the robustness and distinctness of clusters obtained with this value of the resolution parameter using a cross-validation machine learning approach^24, 38^. For each pair of clusters, cells were classified using a multilayer perception classifier. The data was split randomly into five groups and the classifier was trained on 80% of the nuclei, and then repeated five times so that each cell was classified once. This process was repeated 50 times for each cluster pair, resulting in a cluster classification for each nucleus. Nuclei that were classified ambiguously (<40 consistent assignments out of 50 runs) were called “indeterminate” nuclei. The network shown in Extended Data Fig. 3 representing the amount of shared indeterminate cells between clusters was created with the visNetwork package in R.

### Identification of Differentially Expressed Genes Among Myeloid Subgroups

Differentially expressed genes (DEG) were identified for each myeloid subpopulation using the *FindMarkers* function in Seurat with default settings and using the ROC test, which returned the classification power for each marker.

### Functional Annotation of Clusters

Annotation of up-regulated gene lists for each cluster was performed using the DAVID platform, with Gene Ontology using biological process annotation^40, 41^.

### Differences in Relative Abundance of Microglial Subpopulations

We investigated whether the relative abundance of microglial subpopulations differed across PD SN vs NPC SN tissues using Mixed-Effects Association of Single Cells (MASC) analysis^58^. Donor ID, region, and sex were converted to factor variables, and age was scaled as numeric. Region (SN PD vs SN NPC) was assigned as the contrast, donor ID was controlled for as a random effect, and sex and age were controlled for as fixed effects, as designated in the original publication detailing MASC^58^.

MASC was similarly used to investigate the relative abundance of microglial subpopulations across PD tissue regions. PD tissue region (i.e. SN, VTA, SI, and HypoTs) was assigned as the contrast, donor ID and age were controlled for as random effects, and sex was controlled for as a fixed effect.

### Enrichment of PD Susceptibility Genes

Each subpopulation was probed for enrichment of PD GWAS hits using a standard hypergeometric test. The overlap between the PD GWAS gene list described in Nalls et al. 2019^50^ and the differentially expressed gene lists for each cluster (obtained with the Seurat *FindAllMarkers* function) was examined. All p-values were adjusted with the Benjamini-Hochberg correction.

### Immunohistochemistry Validation of Microglial Populations in PD and NDC Tissues

Formalin-fixed paraffin-embedded (FFPE) tissue sections were used, obtained from the contralateral SN from the same donors used for sequencing. To deparaffinize, the sections were dissolved with CitriSolv for three minutes (Decon Labs #. 1601H), followed by two washes in 100% ethanol for one minute each, then 70% ethanol for one minute, and then 3 washes with PBS. To retrieve the antigen epitopes, the sections were microwaved in citrate buffer for 20 minutes followed by three washes with PBS. The sections were then blocked with 5% donkey serum and 0.1% Triton X-100 in PBS. For immunostaining, the sections were incubated overnight at 4°C with the following primary antibodies: TMEM163 (Thermofisher, cat # PA5-114329; 1:100), CD83 (Biolegend, cat # 305302; 1:100), Iba1 (FUJIFILM Wako, cat # 011-27991, 1:500). Primary antibodies were detected using the following appropriate secondary antibodies: Donkey anti-Goat Alexa Fluor 488 (Thermofisher, Cat#: A32814) and Donkey anti-Rabbit Alexa Fluor 555 (Thermofisher, Cat#: A31572). Hoechst 33342 (1μg/ml, Invitrogen, Cat # H3570) was used to counterstain nuclei. After the immunostaining, Trueblack (Biotium, Cat# 23007) was used to reduce the autofluorescence. Sections were mounted in Antifade Mountant (Invitrogen, cat # P36984) under coverslips (No.1.5, VWR) for subsequent microscopy.

### Imaging and Analysis

Confocal images were acquired using Zeiss LSM 900 confocal laser scanning microscope (The Microscopy core of the Columbia Center for Translational Immunology, Columbia University Irving Medical Center) using Plan-Apochromat 20x/0.8 M27 objective with a 1024 X 1024 pixel resolution. A total of 15-20 images were acquired per section and were stored as raw .czi files. Image analysis was performed using the open-access software CellProfiler 4.2.5. Before implementing the automated CellProfiler pipeline-based counting, objects were visualized/compared with a manual count. The raw .czi files were imported into CellProfiler and the corresponding channels were designated based on the metadata of the .czi file. The pipeline steps were as follows: “IdentifyPrimaryObject” command to define the Iba1 positive cells (AF 488 channel), followed by either TMEM163 or CD83 positive objects using the “IdentifyPrimaryObject” command. A robust Background thresholding method was used utilizing the Mean averaging method. To count the CD83+ and Iba1+ double positive cells, as well as the TMEM163+ Iba1+ double positive cells, the RelateObjects command was used. The numerical data were exported using the “ExportToSpreadsheet” command line.

### Validation of Clusters in Other Datasets

The microglia dataset from Siletti et al. 2022^36^, including microglial nuclei from all available brain regions, was downloaded from CellXGene portal. The features were converted from Ensembl IDs to Human gene symbols using BioMart package. This data set was then integrated with microglial nuclei originating from the SN of NPCs from our dataset, using Seurat’s standard integration workflow. Briefly, the raw counts from both datasets were log normalized and then integrated using CCA analysis. The clustering and UMAP visualization were done using 22 PC dimensions and a resolution of 0.4 was selected for further analysis. Region of origin was obtained from the Siletti dataset metadata. As all dissections within the “cerebral nuclei” categorization within the Siletti metadata were from subcortical structures, we have referred to this region as “subcortical structures” accordingly. All other regional labels we have kept unchanged from the Siletti metadata. As described above, MASC was used to investigate the relative abundance of microglial subpopulations across brain regions of origin (tissue region was assigned as the contrast, donor ID was controlled for as a random effect).

### Analysis of ATAC-Seq Data

Input to ATAC-seq pipeline involved selecting cells with single-nucleus ATAC data that also had associated single nucleus RNA-seq data after QC and filtering of the latter. RNA cluster annotations were assigned to the nuclei, and thus preserved in the ATAC-seq data annotation as well. Single nucleus ATAC-seq data was processed and analyzed using the ArchR software package^85^. Quality control for each cell included a minimum TSS score of 4 and a minimum number of 1,000 fragments per cell. We used ArchR’s standard procedure for doublet identification and removal, which includes calculating doublet enrichment score based on simulated doublet enrichment. Standard dimension reductionality and batch effect correction of ATAC data were performed using iterative LSI and Harmony^86^, with “sample” as the batch variable in Harmony.

Peak calling of ATAC-seq data involved first creating pseudobulk replicates using the RNA cluster annotations. Afterward, MACS2 was used for the peak calling step. Wilcoxon rank-sum tests were performed to identify marker peaks for each RNA cluster, while controlling for bias in TSS enrichment and number of fragments. Motif enrichment analyses involved hypergeometric enrichment of a peak annotation for a given motif within up-regulated or down-regulated marker-specific peaks. For enriched motifs, Tn5 insertion location information was leveraged to footprint for transcription factors and determine TF binding locations: ArchR takes into account Tn5 transposase insertion sequence bias for TF footprinting, which would be normalized by subtracting this bias.

Integration of snRNA-seq and snATAC-seq data was performed through the ArchR package as well. Gene scores from the ATAC data were calculated based on accessibility of regulatory elements within 100kb of the gene window. The cross-modality integration in ArchR aligns nuclei from both modalities by comparing the ATAC-seq-derived gene score matrix with the RNA-seq-derived gene expression matrix, constraining cells sharing the same cluster identity.

For creating gene-TF regulatory networks, we leveraged peak-to-gene links to identify peaks that connect to multiple genes, including motifs that may encode for transcription factors and are enriched in cluster-specific peaks. All network figures were created with R package *visNetwork*.

## Acknowledgements

We thank the 33 donors included in this work and their families, whose contribution has allowed us to conduct this research. We thank James Caicedo and Norma Romero for their technical expertise. We thank the New York Brain Bank, the Parkinsonism Brain Bank at Columbia (supported by the Parkinson Foundation USA) for providing all the post-mortem samples used in this study. This project is founded by the Parkinson Foundation Research Center of Excellence at Columbia University. Research reported in this publication was supported by the National Institute of Health (F30AG07461801, R21AG073882, RF1AG058852, R01AG076018, R01NS089674, and T32AI148099, NS107442, NS117583, NS111176, AG064596, UL1TR001873, P30CA013696, P30 AG066462), the DoD (W81XWH-22-1-0127), the Parkinson Foundation (PF-RCE-1948), the Mathers Foundation (MF-2103-01428), ASAP (ASAP-020551), the Ludwig Family Foundation, the Chan Zuckerberg Initiative, and the Thompson Family Foundation. We thank the Columbia University Alzheimer’s Disease Research Center (funded by NIH grant P30AG066462) and the Washington Heights-Inwood Columbia Aging Project (PO1AG07232, R01AG037212, RF1AG054023, R56AG069130, RF1AG066107, UL1TR001873).

## Author Contributions

ZKC, SP, EMB, and VM wrote the manuscript. XE and JPV performed neuropathological characterization of samples. Single nucleus preparations were performed by ZKC and ZP/YZ. Computational analyses were designed and performed by ZKC, AY, JM, and VM. Immunohistochemistry was performed by MR and RT. All authors reviewed the manuscript.

