## Extended Data Figures for "RNA- and ATAC-sequencing Reveals a Unique *CD83+* Microglial Population Focally Depleted in Parkinson’s Disease"

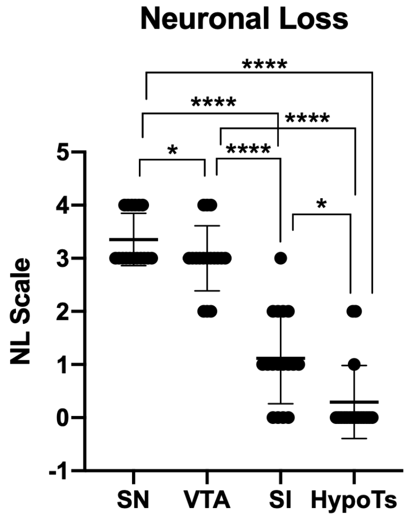


**One Way ANOVA with test for linear trend:**

| R squared (alerting) | R squared (effect size) | F (DFn, DFd) | P value |
|---|---|---|---|
| 0.95 | 0.75 | F (1, 48) = 200.9 | P<0.0001 |

**Extended Data Figure 1. Neuronal loss variation in PD regions.** Neuronal loss (NL) was assessed in tissue regions from each donor, and scored as follows: 0 (no NL), 1 (mild NL), 2 (moderate NL), 3 (severe NL), to 4 (very severe NL). Regions include the substantia nigra (SN), ventral tegmental area (VTA), substantia inominata (SI), and hypothalamus (HypoTs). Each data point indicates the NL score for an individual tissue sample (n=17 per region, donors with missing values/regions were excluded from this analysis). Error bars represent means ± SEM, analyzed by one-way ANOVA $F(2.37, 37.37)= 78.95$, $p<0.0001$ followed by a post-hoc linear test for trend $p<0.0001$, and Tukey's multiple comparisons test: *$p<0.05$, ****$p<0.0001$.

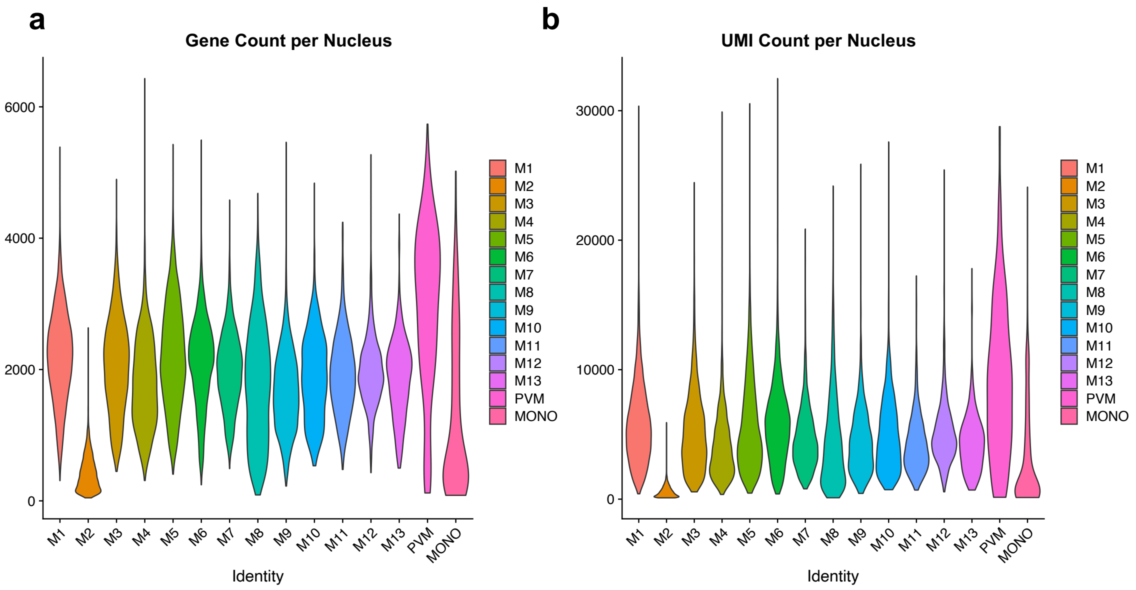


**Extended Data Figure 2. Quality control metrics used for snRNA-seq. a.** Number of unique gene (feature) transcripts expressed for each nucleus sequenced. **b.** Number of total RNA transcripts detected (UMIs) for each nucleus sequenced. Nuclei are categorized by cluster: each column indicates a cluster.

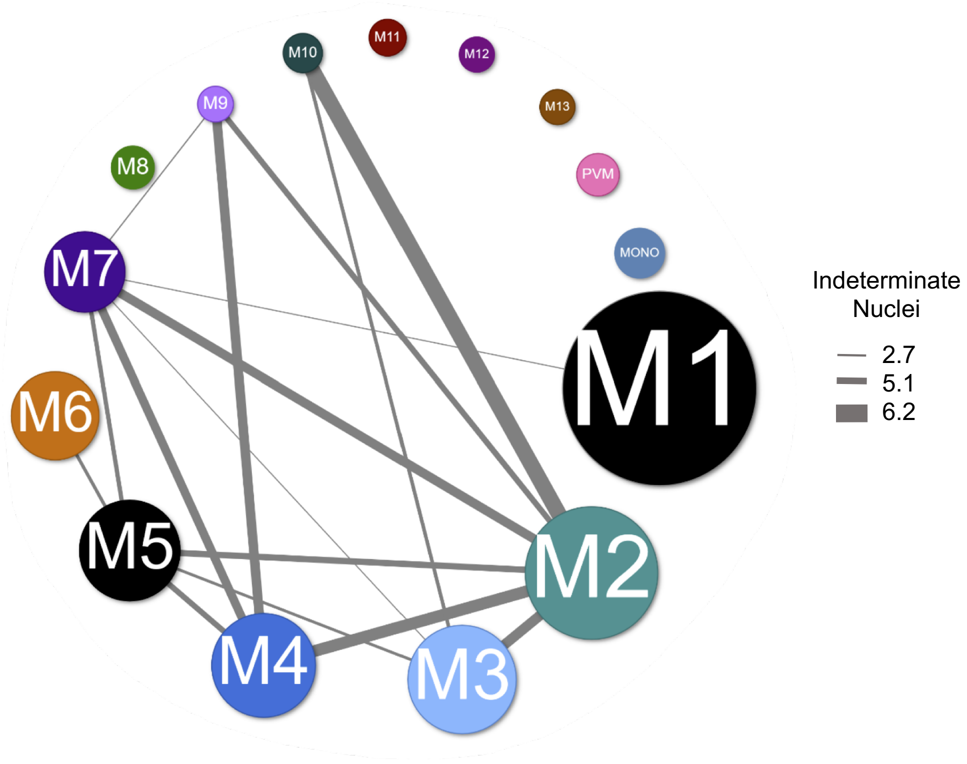


**Extended Data Figure 3. Constellation plot depicting the relationships between different microglial and myeloid clusters.** This analysis was based on a post-hoc machine learning approach using a multilayer perception classifier iterated 50 times. Cells are predicted as indeterminate if they are classified to the same cluster over less than 80% of iterations. Each node represents a cluster, and the edges correspond to the proportion of nuclei that were characterized as indeterminate and shared between clusters. The largest percentage of indeterminate cells between clusters is 6%, suggesting a reliable classification of clusters and cluster stability.

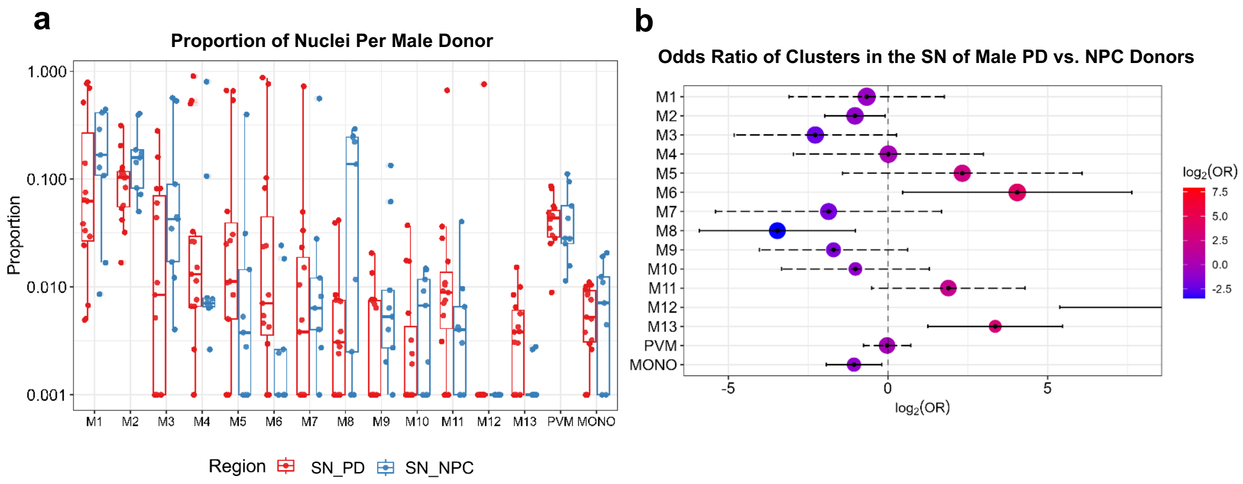


**Extended Data Figure 4. Differential representation of microglial populations in male donors. a.** Boxplot of the proportion of nuclei in each cluster from each individual substantia nigra (SN) tissue sample originating only from male donors within the cohort. Nuclei proportions in each cluster are split by disease condition, Parkinson's disease (SN_PD, red) or non-PD control (SN_NPC, blue). **b.** Abundance of clusters between the SN from male PD and NPC donors using mixed-effects association of single cells (MASC) analysis, controlling for donor, age, and sex as covariates. Forrest plot represents the odds ratio of each nucleus within a given cluster originating from the SN of PD vs NPC donors. Color scale indicates enrichment (red) or depletion (blue) of a cluster in the PD SN compared to NPCs.

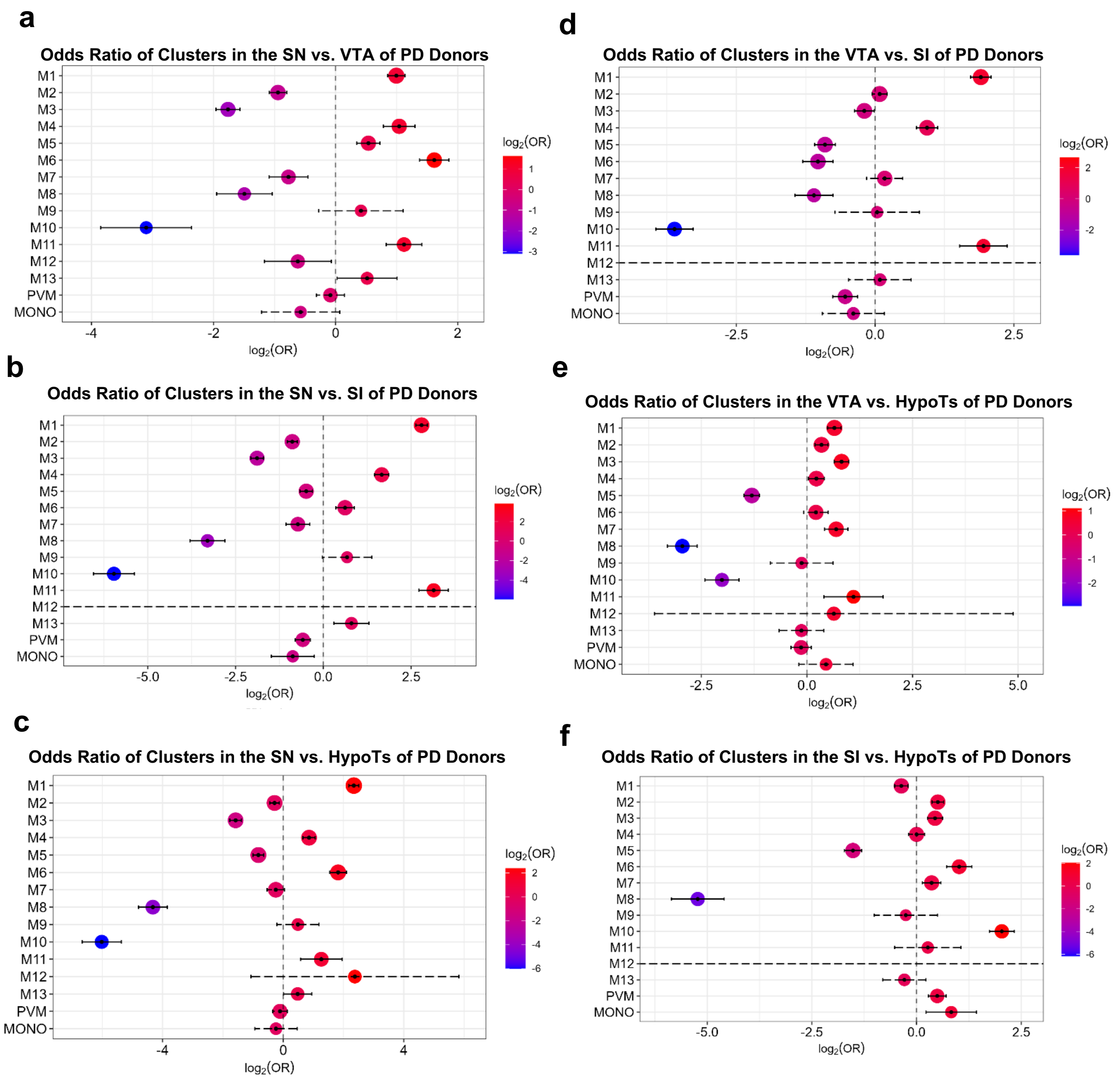


**Extended Data Figure 5. Pairwise comparison of microglial population representation across all Parkinson's disease (PD) donor regions.** The abundance of clusters between all Parkinson's disease (PD) donor regions was assessed using mixed-effects association of single cells (MASC), controlling for donor, age, and sex as covariates**. a.** Forest plot represents the odds ratio of each nucleus within a given cluster originating from the substantia nigra (SN) vs. ventral tegmental area (VTA) of PD donors. **b.** Forest plot represents the odds ratio of each nucleus within a given cluster originating from the SN vs. substantia inominata (SI) of PD donors. **c.** Forest plot represents the odds ratio of each nucleus within a given cluster originating from the SN vs.

rostral hypothalamus (HypoTs) of PD donors. **d.** Forest plot represents the odds ratio of each nucleus within a given cluster originating from the VTA vs SI of PD donors. **e.** Forest plot represents the odds ratio of each nucleus within a given cluster originating from the VTA vs HypoTs of PD donors. **f.** Forest plot represents the odds ratio of each nucleus within a given cluster originating from the SI vs HypoTs of PD donors. Color scale indicates enrichment (red) or depletion (blue) of a cluster in the SN compared to the second indicated region from PD donors.

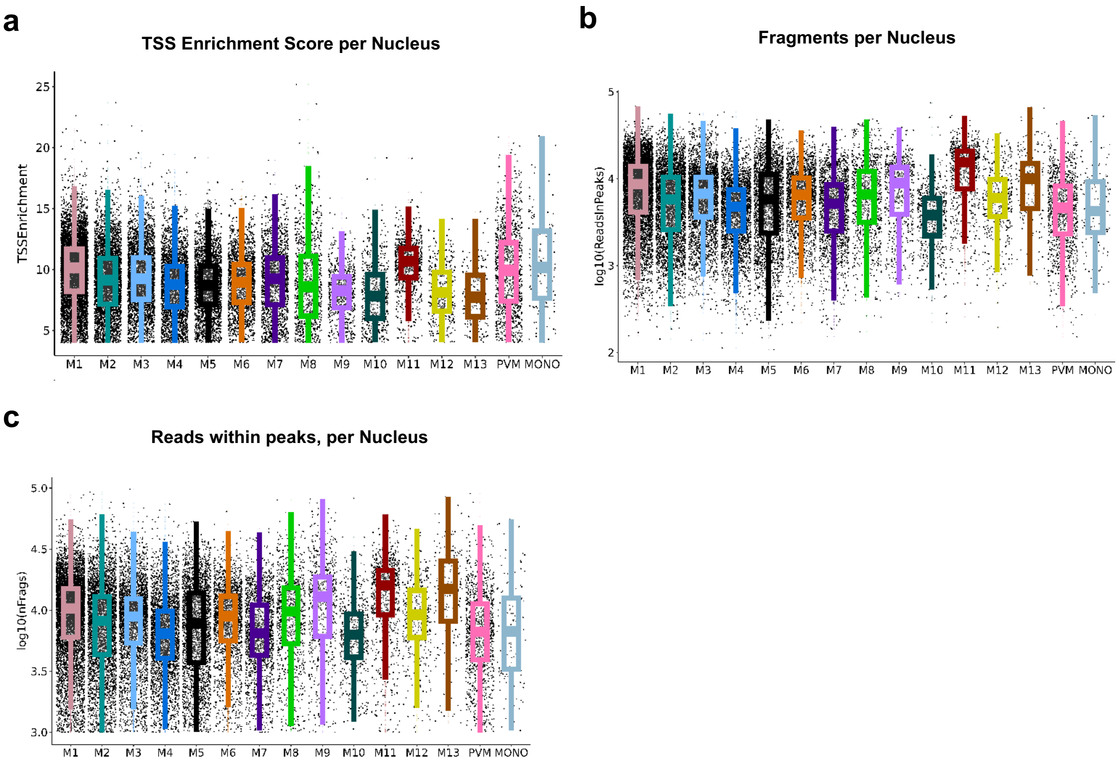


**Extended Data Figure 6. Boxplots of three quality control metrics used for the single nucleus ATAC-sequencing data. a.** Transcription start site (TSS) enrichment score per nucleus. **b.** The number of fragments per nucleus. **c.** Number of reads within peaks per nucleus. Nuclei are categorized by RNA-seq cluster assignment, each column represents an RNA-seq cluster.

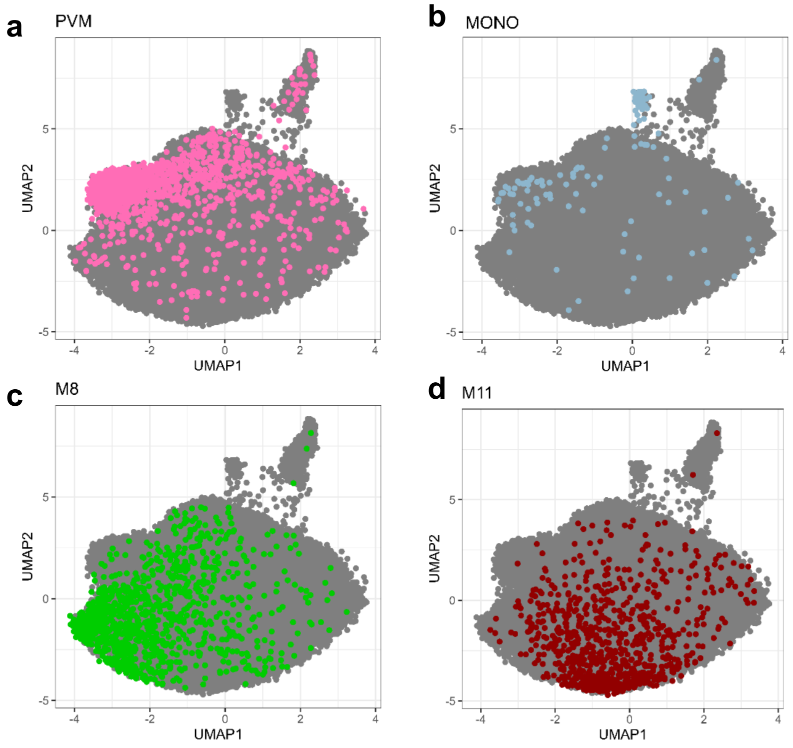


**Extended Data Figure 7**. **ATAC-seq clustering of nuclei from RNA-sequencing clusters PVM, MONO, M8, and M11.** Clustering analysis of single nucleus ATAC-sequencing data performed within the ArchR framework, depicted in UMAP representation. Nuclei are annotated as microglia per their single nucleus RNA-seq cluster label. Nuclei from RNA-seq clusters **a**. PVM, **b**. MONO, **c**. M8, and **d**. M11 are depicted in UMAP space. Each nucleus is represented by a dot. Nuclei from all tissue regions from Parkinson's disease (PD) donors including the substantia nigra (SN), ventral tegmental area (VTA), substantia inominata (SI), and hypothalamus (HypoTs), and the SN from Non-PD controls (NPCs) are included in the UMAP plots.

Extended Data Table 1. Overview of Donor Demographics

Extended Data Table 2. Neuropathological Features of Donors and Tissues Used for Sequencing

Extended Table 3. Differentially expressed genes for each microglial/myeloid cluster, from Seurat *FindAllMarkers* with default settings.

Extended Data Table 4. Differentially expressed genes with classification power for each microglial/myeloid cluster, from Seurat *FindAllMarkers* with ROC test.

Extended Data Table 5. Expression and enrichment (hypergeometric test) for Parkinson's disease GWAS genes for each microglial/myeloid cluster.

Extended Data Table 6. Differential peaks identified for M8.
